# Adiabatic invariants drive rhythmic human motion in variable gravity

**DOI:** 10.1101/674143

**Authors:** N. Boulanger, F. Buisseret, V. Dehouck, F. Dierick, O. White

## Abstract

Natural human movements are stereotyped. They minimise cost functions that include energy, a natural candidate from mechanical and physiological points of view. In time-changing environments, however, motor strategies are modified since energy is no longer conserved. Adiabatic invariants are relevant observables in such cases, although they have not been investigated in human motor control so far. We fill this gap and show that the theory of adiabatic invariants explains how humans move when gravity varies.

---

All living organisms experience a constant terrestrial gravitational acceleration, denoted *as 1g* (9.81 m/*s*^2^). Gravity, “the first thing which you dont think” (A. Einstein), is the most persistent sensory signal in the brain. However, the sensory experiences it generates lack the clear phenomenology of an identifiable stimulus event that characterises sound, sight and even taste. Critically, gravity influences human behaviour more pervasively than any other sensory signal. Exposure to Earth-discrepant gravity – as during spaceflight – leads to dramatic structural and functional changes in the human physiology, including alterations in the cardiovascular [1], neural [2] and musculoskeletal systems [3]. Nowadays the cerebellum appears to be a major structure in gravity perception [4], but we still have no complete understanding of how the brain processes gravity to plan and control actions.

Recent neurocomputational approaches explain behaviour by a mixture of feedback and feedforward mechanisms, conceptualised by internal models [5]: the brain plans an action using available sensory information and makes predictions about the consequences of that action in the environment. Any mismatch between this prediction and the information conveyed by feedback will yield a prediction error used to improve other actions. This mechanism drives motor adaptation. On Earth, gravity is immutable and plays a primary role in minimising prediction errors by providing a strong prior reference.

What is the best way to fundamentally address the role of gravity in motor control? One radical approach consists in challenging the brain by changing a feature of the environment that is never supposed to change: gravity itself. Our original approach is to assess the impact of time-changing gravity on rhythmic biological motion from a purely mechanical vantage point, thereby providing further insights into the fundamental representation of gravity that shapes motor actions. Living organisms are extraordinarily more complex than a simple point-particle body. It is not at all obvious that the actions of a minded human being can be reduced to a standard, simple Lagrangian. Lifting a glass of water off a table requires estimating its weight to adjust the grasping force accordingly. Drinking half of its content with a straw while the glass rests on the table does not, however, allow the brain to program a smaller grasping force, more adapted to the lighter glass [6]. Explicit knowledge of the simplest change in object dynamics is not sufficient to update internal models. Therefore, our working hypothesis is that human actions comply with the behaviour of a mechanical system, even if subject to a slowly changing environment, like a slowly varying gravitational field.

In Mechanics, the most robust way to track the adaptation of a dynamical system to a slow change in the external conditions is through the study of adiabatic invariants and their related action-angle variables describing the system [7]. An adiabatic invariant determines a property of a system that stays approximately constant when external changes occur slowly. Despite their power in revealing constraints on complex dynamical systems, adiabatic invariants have been poorly investigated in biomechanics. For instance, in arm rhythmic motion, the changes in frequency (*df*) occurring during a one-dimensional periodic motion are correlated with changes in energy (*dE*) [8] such that the action variable

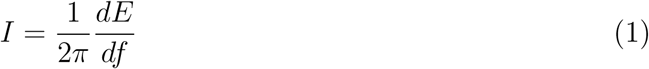

is constant. Action-angle coordinates are usually adopted when the Hamiltonian does not depend explicitly on time. The present work goes beyond previous approaches by immersing participants in a time-dependent gravitational environment where the action variables are not necessarily constant unless the changes in time are adiabatic.

The action-angle variables appeared in the context of classical mechanics in order to study the integrability of dynamical systems with finitely many degrees of freedom. Such systems are said to be *integrable* if the Hamilton-Jacobi equation describing them is completely separable. In the early sixties, the famous Kolmogorov-Arnold-Moser theorem — see [9] for a very interesting book telling the history behind this theorem — brought back the action-angle variables on the scene of classical Mechanics in order to characterise chaotic Hamiltonian systems. Since then and with the seminal works of Nekhoroshev [10, 11] their importance has never faded out. When a Hamiltonian *H*(*P*_*α*_, *Q*^*α*^), *α* = 1,…,*n*, is integrable and leads to bounded trajectories in phase space, action variables may be defined as follows, in terms of a set of phase-space coordinates that separates the Hamiltonian:

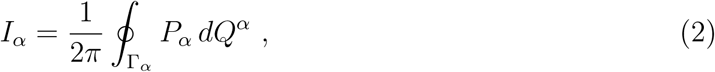

where Γ_*α*_ is the projection of the bounded trajectory in the plane (*P*_*α*_, *Q*^*α*^) for fixed *α*. Once the Hamilton-Jacobi equation is separated in the variables (*Q*^*α*^, *P*_*α*_), on the solution of Hamiltons canonical equations each momentum variable *P*_*α*_ will depend only on its canonically conjugate variable *Q*^*α*^ and on the initial conditions. The action variables give all the conserved quantities of the dynamical system under study, as certified by the Bour-Liouville theorem.

If the Hamiltonian is time-dependent and slowly varying in comparison with the typical period of a cycle, then the action variables are slowly varying too. They are called adiabatic invariants [7, 12, 13] and may be used in a wide range of applications such as in electromagnetism [14], plasma physics [15] and cosmology [16]. Previous works in biomechanics showed the invariance of the action variable when experimental conditions are time-independent [8, 17, 18]. To the best of our knowledge, this concept has never been applied to human motion in time-varying environments. Our approach can reveal the important and otherwise hidden quantities on which the brain relies to plan actions. Advances in this field can potentially not be reached with other, more classical, methods that rest on energy conservation [19]. We therefore designed an experimental set up in which external factors are time-dependent. It is described in the next paragraph.

Six right-handed male participants (40.1 ± 7.2 years old) took part in two centrifugation sessions at QinetiQs Flight Physiological Centre in Linköping, Sweden. The centrifuge was controlled to deliver specific *g*(*t*)-profiles. The real-time control of the orientation of the gondola ensured alignment of local gravity with the long body axis (Fig. 1 inset). One session of centrifugation consisted in a ramp up followed by a ramp down *g*(*t*)-profile for 180s. There were two equivalent sessions separated by a five-minute break bringing the centrifuge back to idle position. The initial 1*g* phases (idle) lasted for 27.4s. Then, the system generated 1.5*g*, 2*g*, 2.5*g*, 3*g*, 2.5*g*, 2*g*, 1.5*g* and 1*g*. Each phase lasted 18.4s and transitions lasted 1.6s (average rate of 0.31*g*/s), except for the first and last ones. We label a given transition by 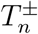 where it is meant that *g*(*t*) goes from the value (*n* + 1)*g*/2 to the value (*n* + 1 + *η*)*g*/2, with *η* = ±1. The increasing (decreasing) gravitational transitions correspond to *η* = +1 (−1). In both cases, *n* ∈ {1, 2, 3, 4}. The first decreasing-*g* series is 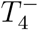 while the last one is 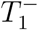 (Fig. 1). A medical flight doctor assessed the participants health status before the experiment. The protocol was reviewed and approved by the Facility Engineer from the Swedish Defence Material Administration (FMV) and an independent medical officer. The experiment was overseen by a qualified medical officer. The study was conducted in accordance with the Declaration of Helsinki (1964). All participants gave informed and written consent prior to the study. A similar protocol was used in a previous study where the human centrifuge is described in detail [20].

**FIG. 1.**
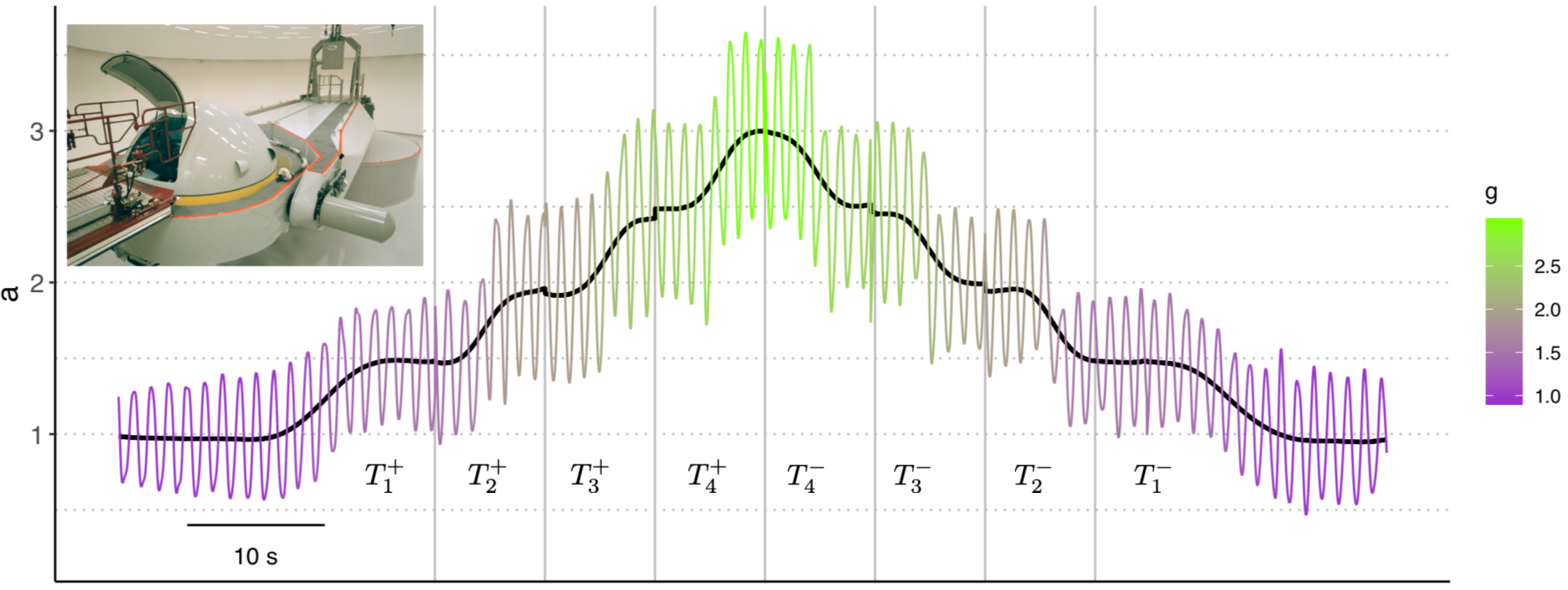
Typical plot of raw data recorded by the accelerometer (coloured line) during a single session of centrifugation (inset). The wireless test object (mass of 0.13 kg) embedded an accelerometer that measured combined gravitational and kinematic accelerations along the objects long axis (AIS326DQ, range 30m/s^2^, accuracy ±0.2m/s^2^). The acceleration signal was sampled at a frequency of 120Hz. The black line depicts local gravity. All accelerations are expressed in units of *g* = 9.81 m/s^2^. The plateau phases are shown for the first and last transitions. For the other transitions, plateau phases and rest periods are not displayed for the sake of clarity but are replaced by vertical lines.

Participants performed upper arm rhythmic movements about the elbow at a free, comfortable pace only during the transitions between gravitational environments, to limit fatigue. When prompted by a GO signal, the participant started to perform the movement while holding an object embedding an accelerometer. The elbow remained in contact with the support. The upper arm produced movements of about 30° with the horizontal. When the operator announced the STOP signal, the participant gently let the object touch the support again while still securing it with his hand. A schematic representation of raw data (acceleration vs time) of one session for one subject is displayed in Fig. 1.

Accelerations *a*(*t*) were numerically integrated and linearly detrended after subtraction of *g*(*t*) to yield the objects speed and position *x*(*t*). The link

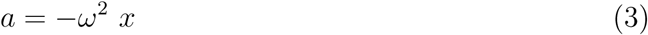

is observed for all participants within a given transition (96 time series): averaged Pearson’s correlation coefficient between *a* and *x* is indeed equal to −0.82 ± 0.1. A typical plot is shown in Fig. 2. In average, *ω* = 6.3 Hz leading to a typical period T= 0.99 s. Hence, we are on safe grounds to assume that the dynamics of the test object along the body axis is compatible with that of a harmonic oscillator, *i.e.*, with a Hamiltonian of the form

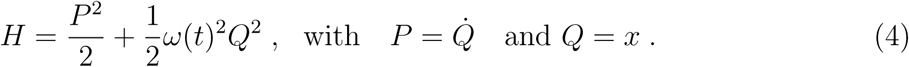

**FIG. 2.**
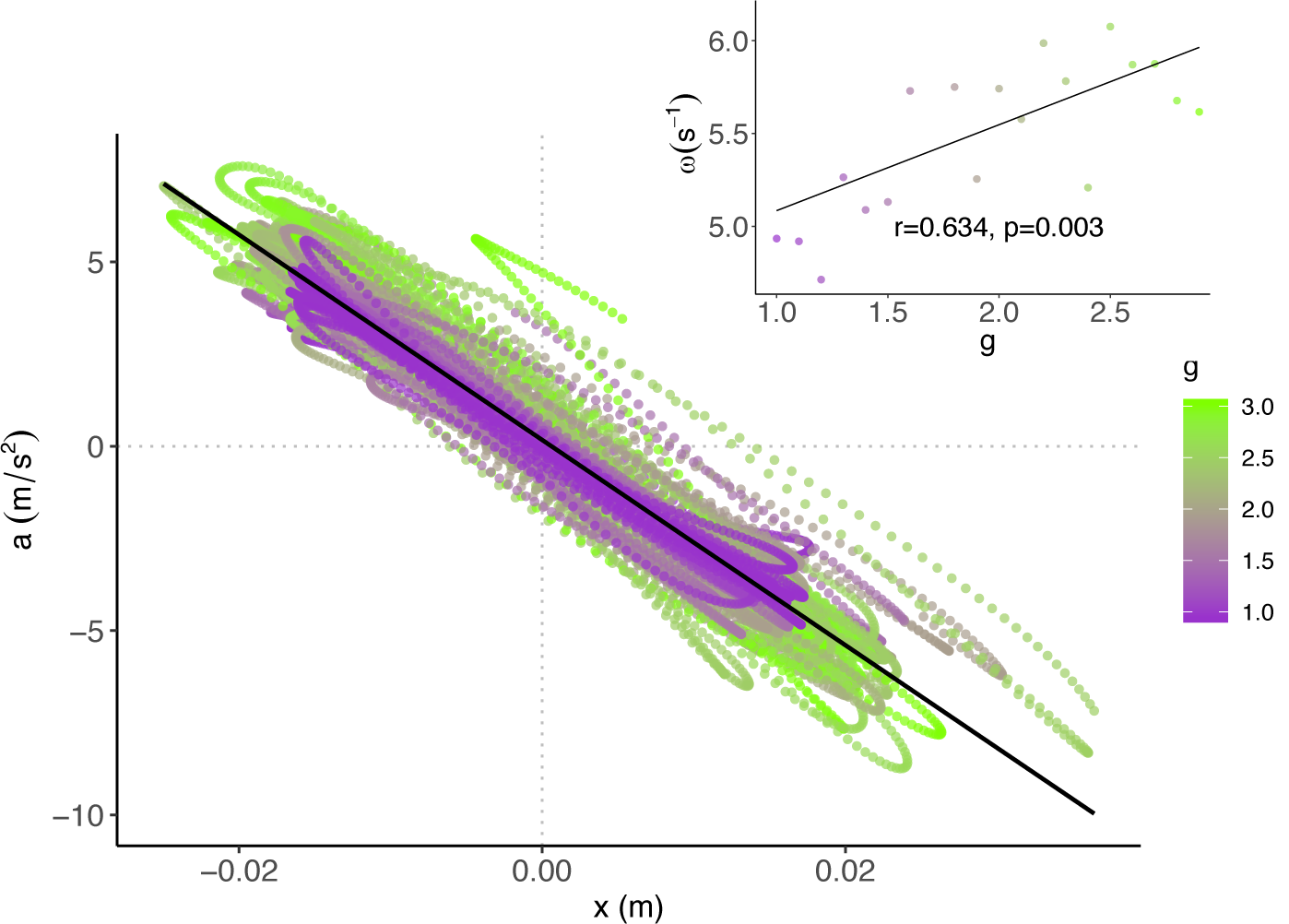
Typical plot of acceleration versus position for the test object during one centrifugation session, same participant as Fig. 1 (coloured points). A global linear regression is shown (solid line). The inset quantifies the significant linear relationship between *ω* and *g*. Dots result from a fit of the form (3) by bins of 0.1 *g*.

Figure 3 depicts a typical phase-space of a complete centrifugation session. Elliptic cycles are clearly visible and are the consequence of the harmonic-oscillator dynamics. The area of these ellipses is slowly changing with *g* as expected from adiabatic invariants theory. Action-angle coordinates (*I, ϕ*) may be defined through the standard definition [7]

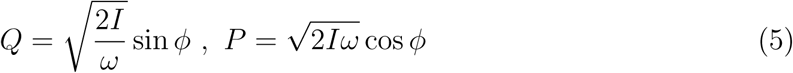

and their equations of motion read

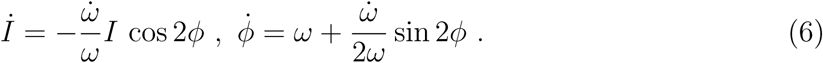

**FIG. 3.**
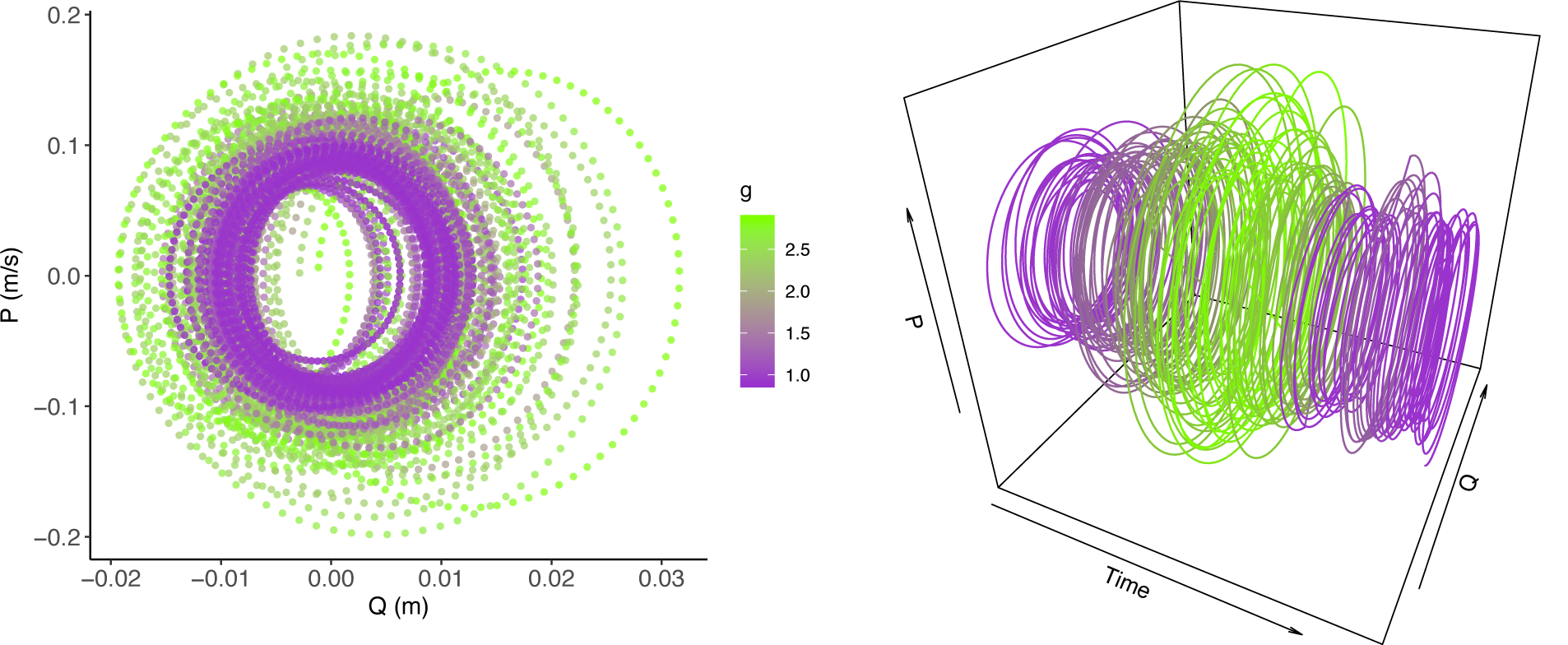
Left panel: Typical phase-space plot of the test object trajectory during one centrifugation session, same participant as Fig. 1. Right panel: Same data but the consecutive cycles are now unfolded along the time dimension.

The parameter of model (4) is the function *ω*(*g*(*t*)). Careful inspection of experimental data let us conclude that *ω*(*g*(*t*)) is compatible with a weakly increasing linear shape, see Fig. 2 inset. Hence we assume

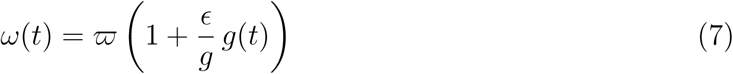

and we will perform computations up to first order in *ϵ* through the rest of the paper. Equation (7) is justified physiologically: muscle stiffness increases with gravitational acceleration to account for the larger motor commands required to perform the same movement. This leads to a modified frequency and *ϵ* > 0. Similarly, muscle stiffness should decrease from normal to microgravity.

Let us now focus on a given transition 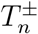. Equation (7) can be adapted to the peculiar shape of *g*(*t*) imposed during the centrifugation:

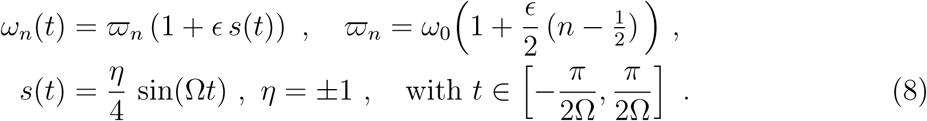

We have shown in [21] that *I*(*t*) and *ϕ*(*t*) can be analytically computed at order *ϵ* from Eq. (6) when *g*(*t*) is of trigonometric form. This gives

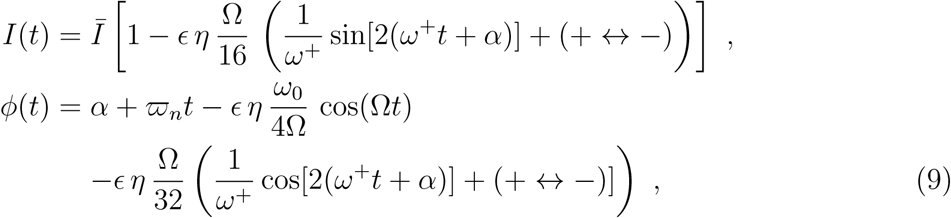

with 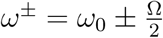 and *ω*_0_ > Ω.

The action variable takes a simpler form when *P* = 0, *i.e.* for *t*_*k*_ such that

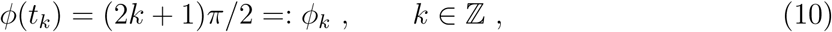

see Eq. (5). The analytical shape of the times *t*_*k*_ such that *ϕ*(*t*_*k*_) = *ϕ*_*k*_ may be complicated but since our goal is the computation of *I*(*t*_*k*_), it is sufficient to work with the lowest order solution 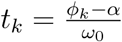, leading to

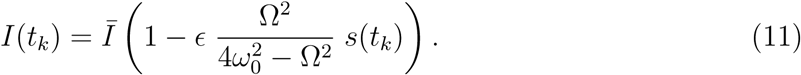

For a given transition 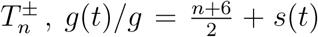. Hence, *I*(*t*_*k*_) = *A*_*n,η*_ + *B g*(*t*_*k*_), where *A*_*n,η*_ and *B* are real constants, and where *B* = *dI/dg* does not depend on *n* and *η*. It allows us to append the transitions and get an affine relation between *I*(*t*_*k*_) and *g*(*t*_*k*_) during the whole centrifugation session:

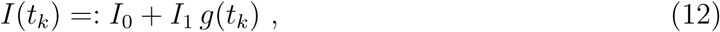

with *I*_0_ ∈ ℝ^+^ and *I*_1_ ∈ ℝ. The shift in *I*(*t*) predicted by Eqs. (9) and (12) extend previous results obtained in Ref. [22] where an analytical shape is obtained for *I*(*t*) with arbitrary *ω*(*t*) provided that the latter is not *C*^∞^.

We have computed phase-space trajectories of all participants in both centrifugation sessions. It is therefore possible to compute the action variable as a function of time. Indeed, Eq. (2) can be rewritten as 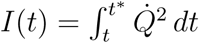, with *t*\* the end of the phase-space cycle starting at *t*. The instant *t*\* > *t* is such that the distance between the points (*Q*(*t*), *P* (*t*)) and (*Q*(*t*\*), *P* (*t*\*)) in phase space is minimal and the difference *t*\* − *t* is as close as possible to T. Once the action variables *I*(*t*) are known, the times *t*_*k*_ such that *P* (*t*_*k*_) = 0 are computed as well as the action variables *I*(*t*_*k*_). Continuous values *I*(*t*_*k*_) of all participants and all trials are finally discretised into 0.1 *g*-bins ranging from 1 to 3 *g*. Each bin contains between 14 and 23 data points. Average values and standard deviations (SD) of *I* normalised to the 1*g* value are finally displayed in Fig. 4.

**FIG. 4.**
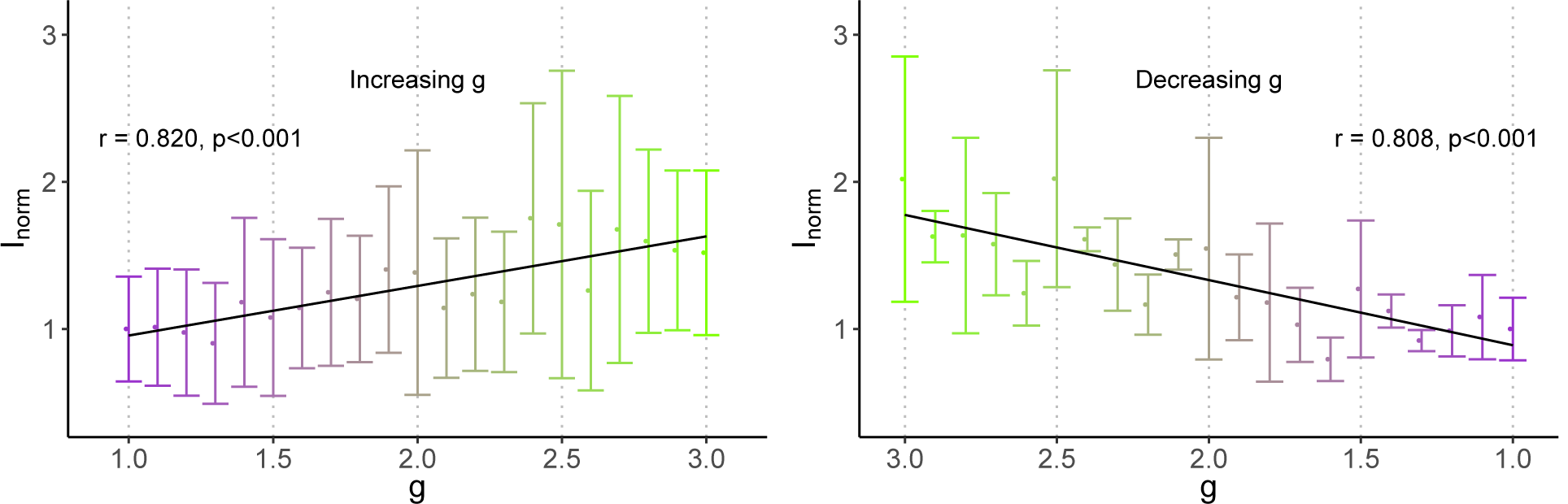
Mean values (and 1 SD error bars) of the adiabatic invariant *I*_norm_ per bin, normalised to the 1*g* value, versus *g*(*t*). Significant linear regressions of the experimental data are depicted as a solid black line together with their Pearsons correlation coefficients and *p*-values. The left panel presents data in the ascending *g*(*t*) phase and the right panel presents data in the descending *g*(*t*) phase. Note that in the descending phase, the horizontal axis is decreasing in order to provide a continuous and chronological reading of the evolution of *I*_norm_.

The adiabatic invariant exhibits a strong and significant positive (*I*_1_ > 0) linear relationship with gravity both in the increasing and decreasing phases (Fig. 4). According to Eq. (1), it shows a higher energetic cost in high gravity for a given change in frequency, which is expected since raising the test object by a height Δ*h* has a potential energetic cost of order *mg*Δ*h*.

Despite this overall coherent dependence of *I* over *g*, we observed asymmetries in the slopes *I*_1_ (Eq. 12) between ascending and descending phases. To quantify this effect, we ran a 2-way repeated measures ANOVA with factors session (1 or 2) and phase (increasing or decreasing). This analysis shows that the slope *I*_1_ is significantly larger in the increasing phase than in the decreasing phase (*I*_1_ = 0.296 ± 0.306 > 0.523 ± 0.219, *p* = 0.037). This asymmetry was not influenced by session (*p* = 0.130). The fact that the adiabatic invariant exhibits a stronger dependence on *g* in the ascending phases is an interesting observation. It indicates that the adiabatic invariant may be modulated more by vestibular and/or proprioceptive gains and re-adjustments of central pattern generators (CPGs) and/or cerebellum activities during that phase. At a spinal cord level indeed, rhythmic movements in mammals are organised by network of interneurons and motor neurons called CPGs [23]. Here, movements of the forearm were generated by CPGs located at cervico-thoracic level [24]. The observation of rapid adaptation of rhythmic forearm movements suggests that vestibular and proprioceptive feedback are the major source of information used by CPGs to ensure adjustments to altered gravity, especially when it increases and becomes more demanding for the control of the task. Interestingly, CPGs receive both inputs from proprioceptive afferents and vestibular pathways. At a supraspinal level, the cerebellum could be the structure integrating the variations of gravity [4], eventually leading to behaviours compatible with adiabatic invariants.

The variability of *I*_norm_ at a given *g* is globally lower in the decreasing than in the increasing-*g* phase as can be seen from the error bars. It suggests habituation takes place because the decreasing-*g* phase always follows the increasing-*g* one. The higher variability during the increasing phase is consistent with the realisation of a movement in a new situation. During the decreasing phase, motor learning achieved in the previous phase made it possible to induce a gradual reduction of variability in order to optimise the movement patterns that are compatible with a simple harmonic oscillator.

In summary, participants show a spontaneous adaptation of their motion that is compatible with the expectation of a simple harmonic oscillator with weakly gravity-dependent frequency. Their adaptation is assessed by the computation of adiabatic invariants, whose experimental behaviour versus *g* comply with our models prediction. We hypothesise that the main biological receptors of time-changing gravity are proprioceptors, such as muscle spindles and Golgi tendon organs that are known to give constant feedback to the CPGs. Adiabatic invariants may thus put realistic constraints on the choices made by spinal and supraspinal nervous structures among an infinite number of possible solutions to a given problem, *i.e.*, the motion of our test object in the present case. Such “hidden” constraints in voluntary motion may be of interest in domains such as rehabilitation and robotics.

Future works might go beyond the harmonic oscillator description of the effective dynamics but still in a phase-space based formalism. As shown in [21], adiabatic invariants can be computed in the case of higher-derivative Hamiltonians of Pais-Uhlenbeck type. Such Hamiltonians could describe rhythmic motions with several frequencies and discrete movements through, *e.g.*, minimal jerk models [25]. We are currently investigating how our model can be generalised by analysing 3D trajectories performed during parabolic flight, therefore also including the very particular case of an absence of gravity [26, 27].

## Acknowledgements

This research was supported by the European Space Agency (ESA) in the framework of the Delta-G Topical Team (4000106291/12/NL/VJ), the “Institut National de la Santé et de la Recherche Médicale” (INSERM) and the “Conseil Général de Bourgogne” (France) and by the Centre National d’Etudes Spatiales grant 4800000665 (CNES). We thank E. Ferrè for inspiring parts of this manuscript

## References

[1] A.E. Aubert et al. Towards human exploration of space: the theseus review series on cardiovascular, respiratory, and renal research priorities. npj Microgravity, 2:16031, 2016.

[2] O. White et al. Towards human exploration of space: the theseus review series on neurophysiology research priorities. npj Microgravity, 2:16023, 2016.

[3] T. Lang, J.J.W.A. Van Loon, S. Bloomfield, L. Vico, A. Chopard, J. Rittweger, A. Kyparos, D. Blottner, I. Vuori, R. Gerzer, and P.R. Cavanagh. Towards human exploration of space: the theseus review series on muscle and bone research priorities. npj Microgravity, 3:8, 2017.

[4] P.R. MacNeilage and S. Glasauer. Gravity perception: The role of the cerebellum. Current Biology, 28, 2018.

[5] M. Kawato. Internal models for motor control and trajectory planning. Curr. Opin. Neurobiol., 9:718–727, 1999.

[6] D.A. Nowak and J. Hermsdrfer. Sensorimotor memory and grip force control: does grip force anticipate a self-produced weight change when drinking with a straw from a cup? Eur. J. Neurosci, 18, 2003.

[7] L. Landau and E. Lifchitz. Physique théorique Tome 1: Mécanique. E. MIR, Moscow, 1988.

[8] M.T. Turvey, K.G. Holt, J. Obusek, et al. Adiabatic transformability hypothesis of human locomotion. Biol. Cybern., 74(107):107–115, 1996.

[9] H.S. Dumas. The KAM story: a friendly introduction to the content, history, and significance of classical Kolmogorov-Arnold-Moser theory. World Scientific, Hackensack, NJ, Apr 2014.

[10] N.N. Nekhoroshev. Behavior of hamiltonian systems close to integrable. Functional Analysis and Its Applications, 5(4):338–339, 1971.

[11] N.N. Nekhoroshev. An exponential estimate of the time of stability of nearly-integrable hamiltonian systems. Uspekhi Matematicheskikh Nauk, 32(6):5–66, 1977.

[12] J. Henrard. The Adiabatic Invariant in Classical Mechanics, pages 60–73. Dessy, 1998.

[13] J.V. Jose and E.J Saletan. Classical dynamics: a contemporary approach. Cambridge Univ. Press, Cambridge, 1998.

[14] J. L. Tennyson, John R. Cary, and D. F. Escande. Change of the adiabatic invariant due to separatrix crossing. Phys. Rev. Lett., 56:2117–2120, May 1986.

[15] J. Notte, J. Fajans, R. Chu, and J. S. Wurtele. Experimental breaking of an adiabatic invariant. Phys. Rev. Lett., 70:3900–3903, Jun 1993.

[16] S. Cotsakis, R. L. Lemmer, and P. G. L. Leach. Adiabatic invariants and mixmaster catastrophes. Phys. Rev. D, 57:4691–4698, Apr 1998.

[17] P.N. Kugler, M.T. Turvey, R.C. Schmidt, and L.D. Rosenblum. Investigating a Nonconservative Invariant of Motion in Coordinated Rhythmic Movements. Ecological Psychology, 2(2):151–189, 1990.

[18] E.E. Kadar, R.C. Schmidt, and M.T. Turvey. Constants underlying frequency changes in biological rhythmic movements. Biol. Cybern., 68:421–430, 1993.

[19] R. McN. Alexander. A minimum energy cost hypothesis for human arm trajectories. Biol. Cybern., 76, 1997.

[20] O. White, J.-L. Thonnard, Ph. Lefèvre, and J. Hermsdörfer. Grip force adjustments reflect prediction of dynamic consequences in varying gravitoinertial fields. Frontiers in Physiology, 9:131, 2018.

[21] N. Boulanger, F. Buisseret, F. Dierick, and O. White. Higher-derivative harmonic oscillators: stability of classical dynamics and adiabatic invariants. Eur. Phys. J., C79(1):60, 2019, 1811.07733.

[22] R.M. Kulsrud. Adiabatic Invariant of the Harmonic Oscillator. Phys. Rev., 106:205–207, 1957.

[23] E. Marder and D. Bucher. Central pattern generators and the control of rhythmic movements. Current Biology, 11, 2012.

[24] E.P. Zehr et al. Possible contributions of cpg activity to the control of rhythmic human arm movement. Can. J. Physiol. Pharmacol., 82, 2004.

[25] T. Flash and N. Hogan. The coordination of arm movements: an experimentally confirmed mathematical model. J. Neurosci, 5, 1985.

[26] O. White et al. Altered gravity highlights central pattern generator mechanisms. J Neurophysiol, 100, 2008.

[27] N. Boulanger, F. Buisseret, V. Dehouck, F. Dierick, and O. White. Rhythmic motion in hyper-and micro-gravity: The role of adiabatic invariants in motor strategy. in preparation, 2019.

